# PLCXD2 preferentially hydrolyzes phosphatidylinositol and supports retinal lipid homeostasis and photoreceptor integrity

**DOI:** 10.64898/2026.09.13.750855

**Authors:** Kaori Kanemaru, Wako Iwata, Akira Katayanagi, Fusui Oka, Yuto Kikuchi, Nozomu Kono, Hyeon-Cheol Lee-Okada, Hideto Osada, Hiroaki Kajiho, Junya Hasegawa, Shin Morioka, Junko Sasaki, Akari Hagiwara, Norimitsu Ban, Junken Aoki, Takehiko Yokomizo, Takehiko Sasaki, Yoshikazu Nakamura

## Abstract

PLCXD proteins have been linked to phosphoinositide metabolism for more than a decade, and PLCXD2 activity has been associated with phosphatidylinositol 4,5-bisphosphate [PI(4,5)P₂] depletion, yet its preferred direct substrate has remained unresolved. Here, cellular lipidomics showed that catalytically active PLCXD2 reduced the major phosphatidylinositol (PI) 38:4 species, with more modest phosphoinositide changes and increased diacylglycerol (DAG) and phosphatidic acid (PA). Using recombinant PLCXD2 and acyl-chain-matched substrates, we found that PLCXD2 hydrolyzed PI substantially more efficiently than PI(4,5)P₂ and most other phosphoinositides tested. In *Plcxd2*-deficient retina, PI abundance was preserved, whereas DAG and PA were reduced without broad changes in major membrane phospholipid classes. PLCXD2 loss was associated with early structural abnormalities at the photoreceptor–bipolar synaptic interface, followed by outer nuclear layer thinning and reduced electroretinographic a-wave responses. These findings identify PLCXD2 as a PI-preferring mammalian PLC and reveal an endogenous role in retinal lipid homeostasis and outer retinal integrity.

## Introduction

Phospholipase C (PLC) enzymes catalyze the cleavage of phospholipids to generate diacylglycerol (DAG) and phosphorylated headgroups, but their substrate usage differs markedly across biological systems. Canonical mammalian PLCs are best known for hydrolyzing phosphatidylinositol 4,5-bisphosphate [PI(4,5)P₂] to generate DAG and inositol 1,4,5-trisphosphate, thereby coupling membrane signaling to protein kinase C activation and intracellular Ca²⁺ mobilization^1–5^. PI-directed PLC activity is also well established in bacteria, where PI-specific PLCs directly hydrolyze unphosphorylated phosphatidylinositol (PI)^6–9^. In mammalian phosphoinositide signaling, however, unphosphorylated PI is generally regarded as the upstream precursor from which phosphorylated phosphoinositides are generated^10–13^. Because PI constitutes a substantially larger membrane lipid pool than individual phosphoinositides, direct access to PI by a mammalian PLC could have consequences distinct from conventional PI(4,5)P₂ hydrolysis, potentially affecting both DAG production and the availability of precursor lipid for phosphoinositide synthesis.

PLC X-domain-containing proteins (PLCXDs) form an atypical branch of the mammalian PLC family. Initial characterization of PLCXD proteins showed that their expression increased cellular inositol phosphate turnover, supporting phospholipase activity but without directly identifying the phospholipid substrate responsible for this activity^14^. Subsequent studies increasingly linked PLCXD proteins to PI(4,5)P₂ homeostasis. In *Drosophila*, catalytic activity of dPLCXD is required for a phosphatase-independent PTEN pathway that limits PI(4,5)P₂ accumulation on endosomes^15^. More recently, mammalian PLCXD2 was shown to be a constitutively active PLC whose restraint by the postsynaptic receptor GPR158 prevents excessive PI(4,5)P₂ depletion and supports neuronal membrane organization^16^. However, these cellular studies did not establish whether PI(4,5)P₂ is the preferred direct substrate of PLCXD2. Thus, despite increasing evidence linking PLCXD2 activity to PI(4,5)P₂ homeostasis, its direct substrate preference remained unresolved. Very recently, another mammalian PLCXD family member, PLCXD1, was reported to catalyze the conversion of PI to DAG, with its catalytic activity required for insulin-stimulated lipogenesis^17^.

For PLCXD2 itself, a central unresolved question remained: which lipid is its preferred direct substrate, particularly whether PLCXD2 primarily acts on PI(4,5)P₂ or unphosphorylated PI. Cellular depletion of PI(4,5)P₂ associated with PLCXD2 activity is consistent with direct PI(4,5)P₂ hydrolysis, but does not by itself establish PI(4,5)P₂ as the predominant direct substrate. Alternatively, PLCXD2-mediated hydrolysis of the upstream PI pool could influence PI(4,5)P₂ and other phosphoinositides indirectly by altering precursor availability. Distinguishing between these possibilities requires direct comparison of PLCXD2 activity toward PI and phosphorylated phosphoinositides under matched biochemical conditions. This distinction is important because PI and PI(4,5)P₂ occupy distinct positions in membrane lipid metabolism: PI is the precursor for phosphoinositide synthesis and a substantially larger lipid pool, whereas PI(4,5)P₂ is a low-abundance regulatory lipid^10,11^. Preferential PI hydrolysis would therefore position PLCXD2 to influence both phosphoinositide precursor availability and DAG-linked lipid metabolism. It is also important to determine how PLCXD2 catalytic activity affects cellular PI-linked lipid metabolism and whether endogenous PLCXD2 contributes to lipid homeostasis *in vivo*.

Here, we define the lipid substrate preference of PLCXD2 and examine its cellular and physiological consequences. In HEK293 cells, expression of catalytically active PLCXD2 caused a pronounced reduction in the major PI 38:4 species, accompanied by more modest changes in phosphoinositide pools and marked increases in DAG and phosphatidic acid (PA). Direct biochemical comparison using recombinant PLCXD2 and acyl-chain-matched lipid substrates showed that PLCXD2 hydrolyzed PI substantially more efficiently than PI(4,5)P₂ and most other phosphoinositides tested. *In vivo*, genetic loss of PLCXD2 reduced retinal DAG and PA without broadly altering major membrane phospholipid classes. *Plcxd2* deficiency was additionally associated with early abnormalities at the photoreceptor–bipolar synaptic interface, followed by photoreceptor loss and impaired photoreceptor responses. Together, these findings establish a pronounced preference of PLCXD2 for PI over PI(4,5)P₂, link its catalytic activity to cellular PI-linked lipid metabolism, and reveal an endogenous role for PLCXD2 in retinal DAG–PA homeostasis and outer retinal integrity.

## Results

### PLCXD2 activity alters PI-linked lipid metabolism

Previous studies linked PLCXD2 activity to PI(4,5)P₂ depletion and PI(4,5)P₂-dependent cellular phenotypes^15,16^. We therefore first asked whether PLCXD2 catalytic activity predominantly perturbs PI(4,5)P₂ itself or produces a broader pattern of changes across PI-linked lipid metabolism. FLAG-tagged wild-type PLCXD2 (PLCXD2 WT) or a catalytically inactive mutant (PLCXD2 MT) was expressed in HEK293 cells (Fig. 1a), and cellular phosphoinositides were quantified by LC-MS/MS-based lipidomics. The catalytically inactive H57L/H132L mutant, previously used to disrupt PLCXD2 phospholipase activity^16^, allowed lipid changes dependent on PLCXD2 catalytic activity to be distinguished from effects associated with protein expression itself.

**Figure 1.**
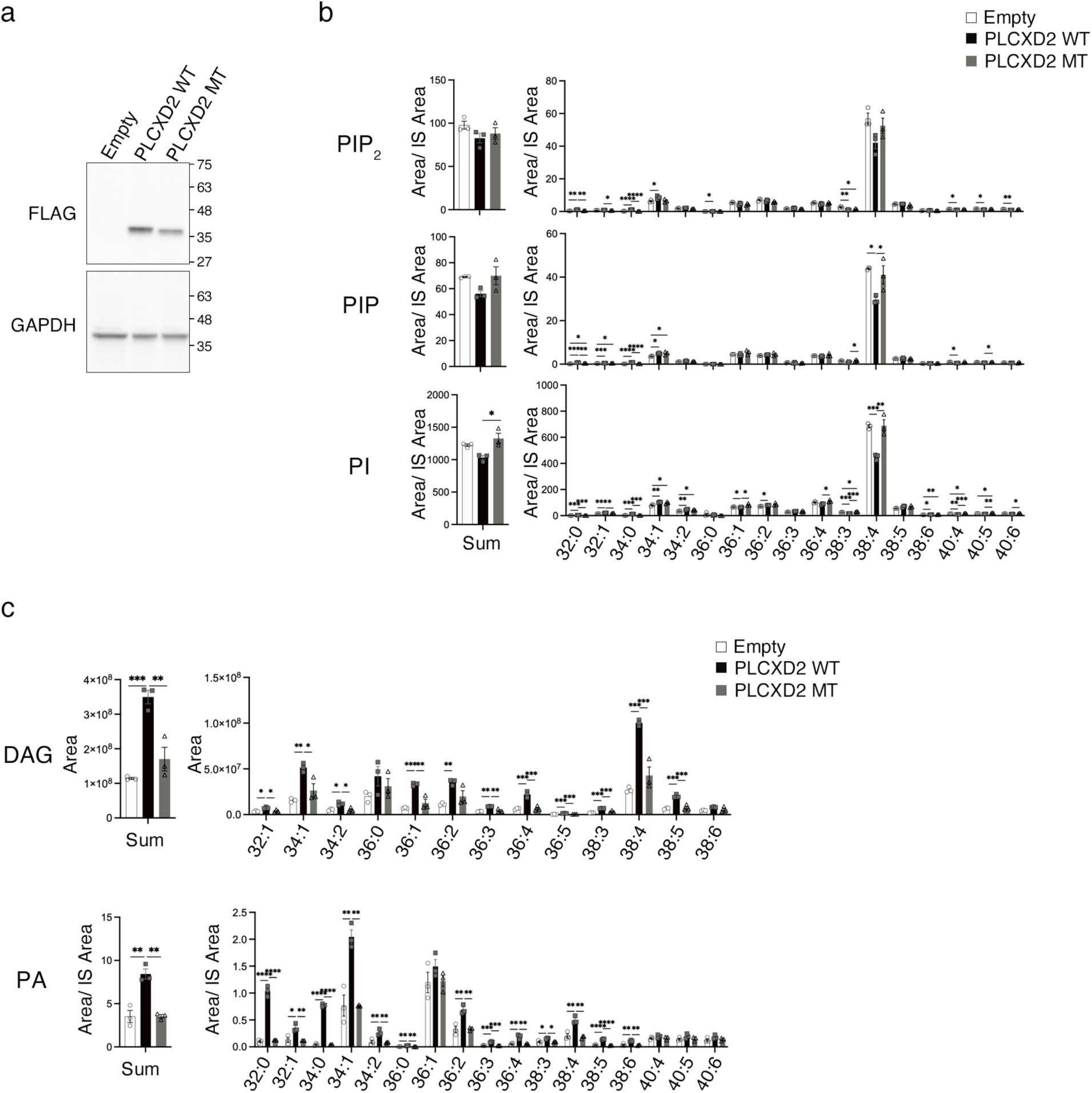
PLCXD2 activity alters PI-linked lipid metabolism. (a) Expression of FLAG-tagged PLCXD2 WT and catalytically inactive PLCXD2 MT in HEK293 cells. FLAG and GAPDH were detected by immunoblotting. (b) LC–MS/MS analysis of PIP₂, PIP, and PI extracted from HEK293 cells expressing empty vector, PLCXD2 WT, or PLCXD2 MT. The left panels show the sum of the normalized signals for all detected molecular species (Sum), and the right panels show individual molecular species. (c) LC–MS/MS analysis of DAG and PA in HEK293 cells expressing empty vector, PLCXD2 WT, or PLCXD2 MT. The left panels show the summed signals for all detected molecular species (Sum), and the right panels show individual molecular species. PIP₂, PIP, PI, and PA levels were calculated as the analyte peak area divided by the corresponding internal-standard (IS) peak area. DAG levels are presented as LC–MS/MS peak areas normalized to 1 × 10^5^ cells. Data are presented as mean ± SD from three biologically independent experiments. Statistical significance was assessed by one-way ANOVA followed by Tukey’s multiple-comparisons test. *P < 0.05, **P < 0.01, ***P < 0.001, and ****P < 0.0001.

The LC-MS/MS method used here quantified mono- and bis-phosphorylated PI species without resolving positional phosphate isomers; these pools are therefore referred to as PIP and PIP₂, respectively. Because PI(4)P and PI(4,5)P₂ represent the predominant cellular PIP and PIP₂ species, respectively, these measurements are expected to be weighted toward these abundant isomers, although contributions from other positional isomers cannot be excluded.

PLCXD2 WT expression caused a modest decrease in total PIP₂ relative to empty-vector control and PLCXD2 MT-expressing cells, although the difference did not reach statistical significance (Fig. 1b). Analysis of individual molecular species revealed a similar downward trend in PIP₂ 38:4, the predominant PIP₂ species in these cells. In the PIP pool, PIP 38:4 was significantly reduced upon PLCXD2 WT expression (Fig. 1b). Comparable decreases were not broadly observed among other PIP and PIP₂ species, indicating that the phosphoinositide changes in HEK293 cells were concentrated in specific molecular species rather than reflecting uniform depletion of the entire PIP or PIP₂ pool.

We next examined PI, the common precursor of all phosphoinositides. Total PI abundance was only modestly altered, whereas molecular-species analysis revealed a pronounced reduction in PI 38:4 in HEK293 cells (Fig. 1b). In contrast, a comparable decrease was not observed across other PI species. Because phosphorylation of the inositol headgroup does not alter the acyl-chain composition of PI, the reduction in PI 38:4, together with the decrease in PIP 38:4 and the downward trend in PIP₂ 38:4 raised the possibility that PLCXD2 perturbs the PI pool upstream of phosphoinositide synthesis.

PI is substantially more abundant than PI(4,5)P₂ in mammalian cells^10,11^. Thus, if PLCXD2 can access the cellular PI pool, even partial turnover of PI could have a considerably greater impact on DAG production and downstream glycerolipid metabolism than hydrolysis of the much smaller PI(4,5)P₂ pool. We therefore asked whether the reduction in PI associated with PLCXD2 activity was accompanied by accumulation of lipids downstream of PLC-mediated PI hydrolysis. We quantified DAG, the immediate product of PLC-catalyzed phosphodiester bond cleavage, and PA, which can be generated from DAG by diacylglycerol kinases^18,19^.

PLCXD2 WT expression markedly increased total DAG compared with both empty-vector control and PLCXD2 MT-expressing cells (Fig. 1c). Among individual species, DAG 38:4 showed a particularly large increase in signal intensity, consistent with the reduction in the major PI 38:4 pool observed in these cells. However, DAG increases were not restricted to 38:4 but extended across multiple molecular species. PA was likewise increased in PLCXD2 WT-expressing cells and showed changes across multiple molecular species (Fig. 1c). Together, the broader changes in DAG and PA suggest that the metabolic consequences of PLCXD2 activity involve additional changes in downstream glycerolipid metabolism beyond those reflected by the reciprocal changes in PI 38:4 and DAG. By contrast, major membrane phospholipid classes showed only limited and heterogeneous changes, without a consistent pattern across lipid classes (Supplementary Fig. 1).

Together, these cellular lipidomic data revealed a pattern not dominated by uniform PI(4,5)P₂ depletion. Instead, a pronounced decrease in PI 38:4 was accompanied by more modest, 38:4-centered changes in PIP and PIP₂, together with marked accumulation of DAG and PA. These observations were difficult to reconcile with a model in which PLCXD2 acts solely as a PI(4,5)P₂-directed PLC and raised the possibility that unphosphorylated PI itself is a major direct substrate of PLCXD2.

### Direct biochemical comparison identifies PI as the preferred substrate of PLCXD2

To discriminate directly between PI(4,5)P₂-directed and PI-directed models of PLCXD2 activity, we examined the substrate preference of recombinant PLCXD2 using defined phospholipid substrates. Recombinant PLCXD2 WT and the catalytically inactive MT protein were prepared in a cell-free expression system and subjected to *in vitro* phospholipase assays (Fig. 2a). To relate the biochemical analysis to the lipid species prominently affected in HEK293 cells, we used PI 38:4 and the corresponding phosphoinositide species with a 38:4 sum composition as substrates.

**Figure 2.**
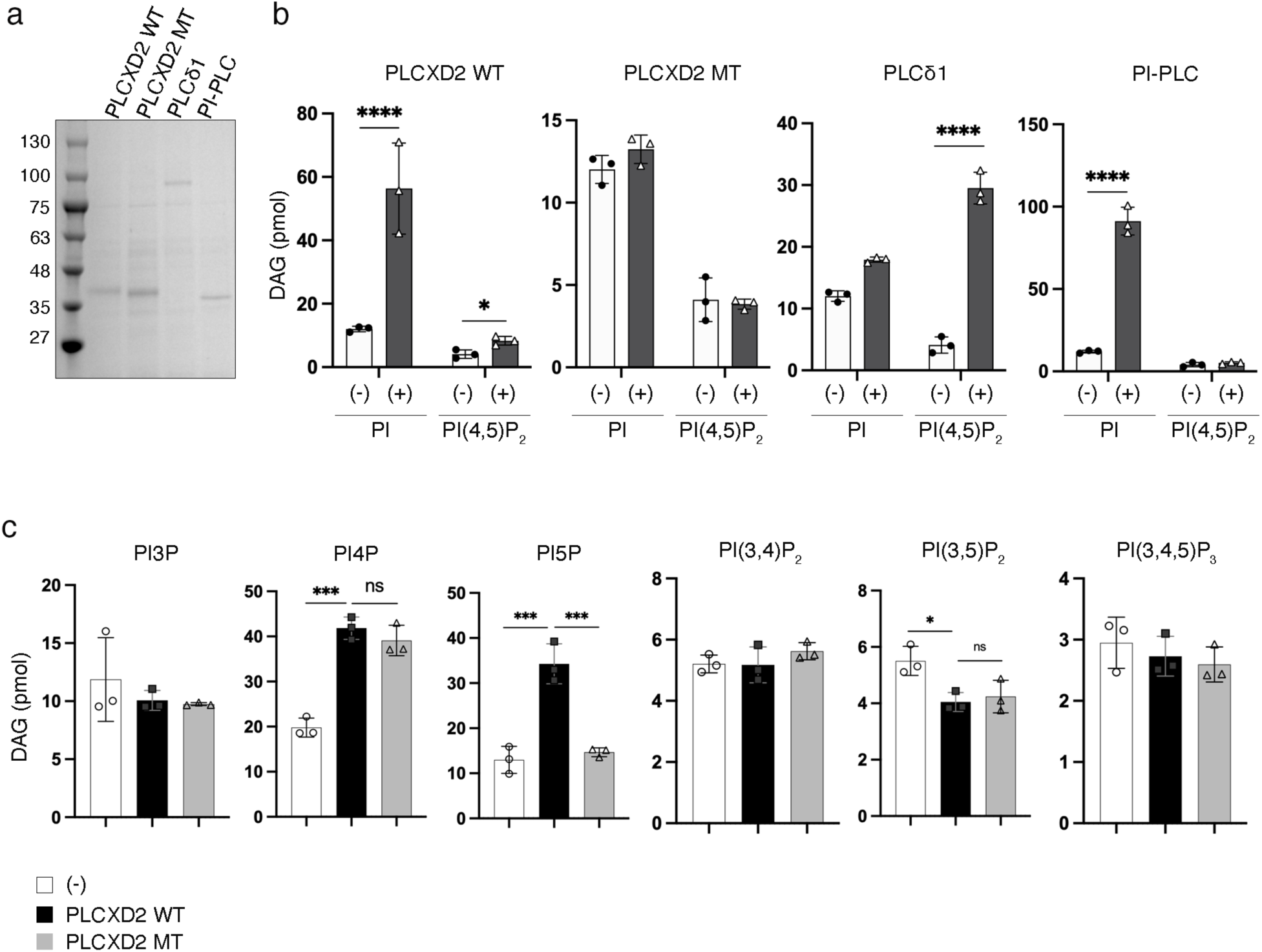
PLCXD2 preferentially hydrolyzes phosphatidylinositol *in vitro*. (a) SDS–PAGE and Coomassie Brilliant Blue staining of recombinant PLCXD2 WT and PLCXD2 MT preparations used for the biochemical assays, together with mammalian PLCδ1 and bacterial PI-PLC reference enzymes. (b) DAG production from matched PI 38:4 or PI(4,5)P₂ 38:4 substrates in reactions containing the indicated recombinant enzymes. PLCδ1 and bacterial PI-PLC served as reference enzymes for PI(4,5)P₂-directed and PI-directed PLC activity, respectively. DAG production was quantified by LC–MS/MS. For each substrate, the no-enzyme condition was shared across the enzyme comparisons and assayed in parallel; the same no-enzyme values are therefore shown in the corresponding graphs. (c) DAG production from PI3P, PI4P, PI5P, PI(3,4)P₂, PI(3,5)P₂, and PI(3,4,5)P₃ in reactions containing no enzyme, PLCXD2 WT, or PLCXD2 MT. In (b) and (c), (−) denotes reactions without enzyme; in (b), (+) denotes reactions containing the recombinant enzyme indicated above each graph. Data are presented as mean ± SD from three independent assays. Statistical significance was assessed by one-way ANOVA followed by Dunnett’s multiple-comparisons test in (b) and Tukey’s multiple-comparisons test in (c). *P < 0.05, **P < 0.01, ***P < 0.001 and ****P < 0.0001.

Phospholipase activity was quantified by measuring DAG production using LC-MS/MS. As reference enzymes, mammalian PLCδ1 efficiently generated DAG from PI(4,5)P₂, whereas bacterial PI-PLC strongly hydrolyzed unphosphorylated PI (Fig. 2b), confirming that the assay could distinguish PI(4,5)P₂-directed from PI-directed PLC activity. PLCXD2 WT generated substantial amounts of DAG from PI, whereas the catalytically inactive PLCXD2 MT showed no corresponding increase (Fig. 2b), demonstrating catalytic activity-dependent PI hydrolysis by recombinant PLCXD2. PLCXD2 WT also hydrolyzed PI(4,5)P₂, but to a much lesser extent. Under the same assay conditions, background-subtracted DAG production from PI was approximately an order of magnitude greater than that from PI(4,5)P₂. Thus, in contrast to the PI(4,5)P₂ preference of canonical mammalian PLCs, PLCXD2 displayed a pronounced preference for unphosphorylated PI.

We next examined whether PLCXD2 could directly hydrolyze other phosphorylated phosphoinositides. Under the same assay conditions, no clear PLCXD2 catalytic activity-dependent DAG production was detected from PI3P, PI(3,4)P₂, PI(3,5)P₂, or PI(3,4,5)P₃ (Fig. 2c). PI4P-containing reactions showed elevated DAG levels, but similar increases were observed with PLCXD2 WT and MT, arguing against PLCXD2-dependent PI4P hydrolysis. By contrast, PI5P supported a significant WT-dependent increase in DAG, indicating that PI5P can also serve as a PLCXD2 substrate *in vitro*. Together, these matched biochemical analyses demonstrate that PLCXD2 directly hydrolyzes PI and does so substantially more efficiently than PI(4,5)P₂ and the other phosphoinositides tested, establishing PI as its preferred substrate under these assay conditions.

### Endogenous PLCXD2 contributes to retinal DAG and PA pools *in vivo*

We next asked whether endogenous PLCXD2 contributes to PI-linked lipid metabolism *in vivo*. To this end, we generated *Plcxd2* knockout mice using CRISPR/Cas9-mediated genome editing. Two guide RNAs flanking exons 2 and 3 were used to generate a deletion encompassing these exons (Fig. 3a). The wild-type and mutant alleles were distinguished by genomic PCR using the F1, R1, and R2 primers indicated in Fig. 3a (Fig. 3b). Analysis of *Plcxd2* cDNA isolated from knockout retina showed direct joining of exon 1 to exon 4, which was confirmed by sequencing of the exon junction (Fig. 3c). This deletion introduces a frameshift followed by a premature termination codon at amino acid 64, predicting a severely truncated protein lacking the catalytic X domain. The mutant allele is therefore expected to abolish PLCXD2 phospholipase activity.

**Figure 3.**
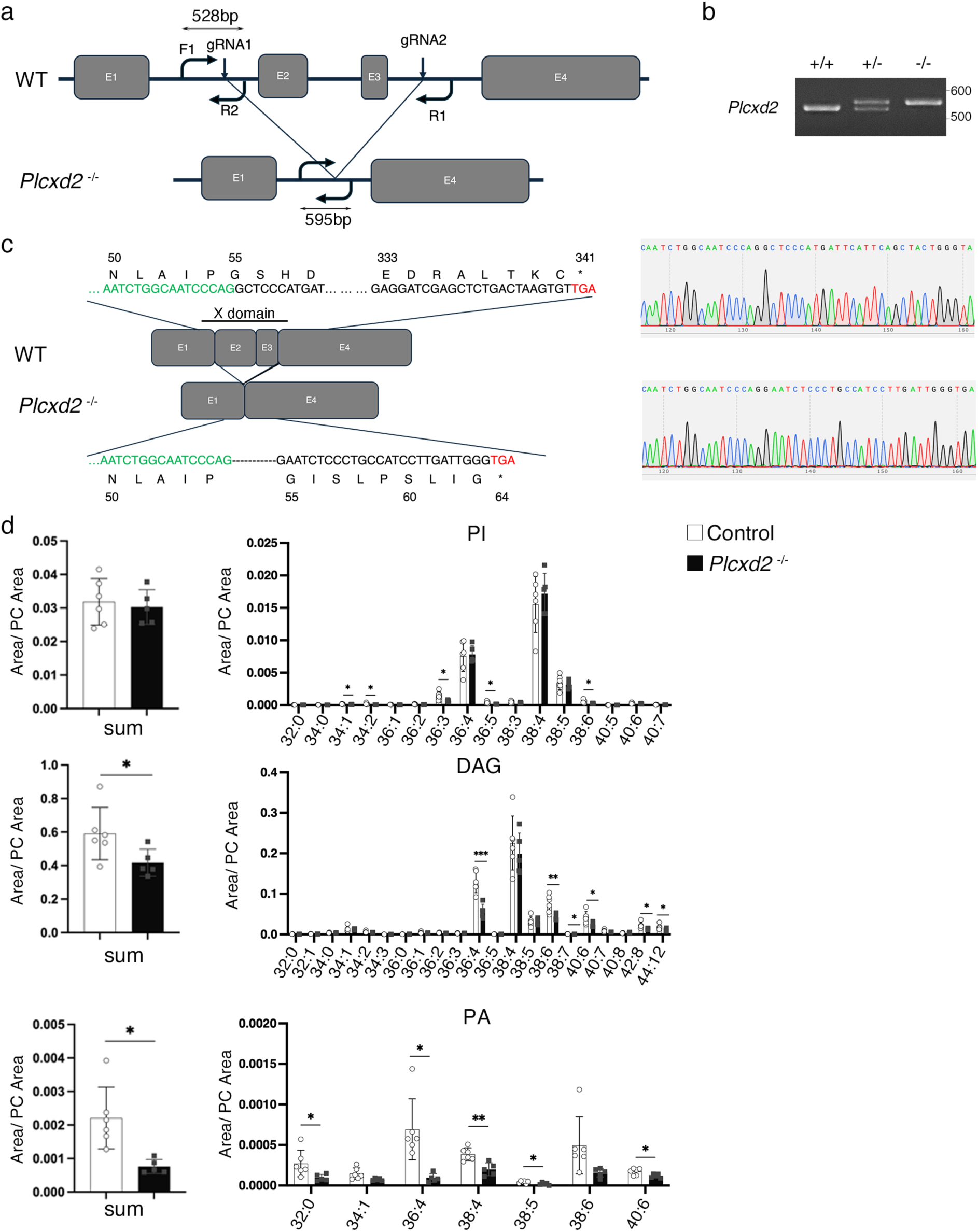
Endogenous PLCXD2 contributes to retinal DAG and PA pools in vivo. (a) CRISPR/Cas9 strategy used to generate the Plcxd2 knockout allele. Exons are indicated as E1–E4. Guide RNAs flanking exons 2 and 3 generated a deletion encompassing both exons. The positions of the genotyping primers F1, R1, and R2 and the expected PCR product sizes for the wild-type and deletion alleles are indicated. (b) Representative genomic PCR analysis of wild-type (+/+), heterozygous (+/−), and homozygous knockout (−/−) mice using the primers shown in (a). The wild-type and deletion alleles generated products of approximately 528 and 595 bp, respectively. (c) Retinal cDNA analysis. The knockout transcript joins exon 1 directly to exon 4, resulting in a frameshift and predicted premature termination at amino acid 64, upstream of the catalytic X domain. Representative Sanger sequencing chromatograms confirm the exon 1–exon 4 junction. (d) LC–MS/MS analysis of PI, PA, and DAG in retinas from 21-week-old male control and *Plcxd2* knockout mice. The sum of the PC-normalized signals for all detected molecular species (Sum) and the signals for individual molecular species are shown. Data are presented as mean ± SD (control, n = 6 mice; knockout, n = 5 mice). Statistical significance was assessed using two-tailed Welch’s t-test. *P < 0.05, **P < 0.01, and ***P < 0.001.

We then examined retinal lipid composition by LC-MS/MS. Because the small size of the retina made normalization by tissue weight unreliable, lipid abundance was normalized to total phosphatidylcholine (PC) within each sample, an abundant membrane phospholipid that could be robustly quantified across samples. We first examined whether loss of PLCXD2 altered steady-state retinal PI abundance. However, neither total PI nor PI 38:4, the molecular species most prominently decreased by PLCXD2 overexpression in HEK293 cells, was increased in *Plcxd2* knockout retina (Fig. 3d). Thus, steady-state retinal PI abundance was maintained despite PLCXD2 deficiency.

In contrast, total DAG was significantly reduced in *Plcxd2* knockout retina (Fig. 3d). The decrease extended across several DAG molecular species, whereas the predominant DAG 38:4 species was largely preserved. Total PA was also markedly reduced, with decreases observed across multiple PA molecular species (Fig. 3d). Thus, although loss of PLCXD2 did not result in detectable accumulation of PI, the reduction in DAG is consistent with a contribution of endogenous PLCXD2 to retinal DAG generation, accompanied by associated changes in the PA pool.

The decreases in DAG and PA were not accompanied by comparable changes in the total abundance of other major membrane phospholipid classes, including PE, PG, PS, and SM (Supplementary Fig. 2). Thus, the reduction in DAG and PA did not reflect a generalized loss or disruption of retinal membrane lipids. These findings show that endogenous PLCXD2 contributes to retinal DAG–PA pools without broadly perturbing membrane lipid composition. Although the retinal data do not identify the lipid substrate consumed by PLCXD2 *in vivo*, they are consistent with the biochemical and cellular evidence that PLCXD2 feeds PI-linked lipid metabolism into the DAG–PA axis.

### PLCXD2 deficiency causes early outer retinal abnormalities followed by photoreceptor degeneration

We next examined whether PLCXD2 deficiency affects retinal organization and function. At 3 months of age, retinal lamination was grossly preserved in *Plcxd2* knockout mice, and quantitative analysis revealed no significant differences in the thickness of the outer nuclear layer (ONL), outer plexiform layer (OPL), inner nuclear layer (INL), or inner plexiform layer (IPL) compared with control mice (Fig. 4a). Thus, overt retinal degeneration was not yet apparent at this stage.

**Figure 4.**
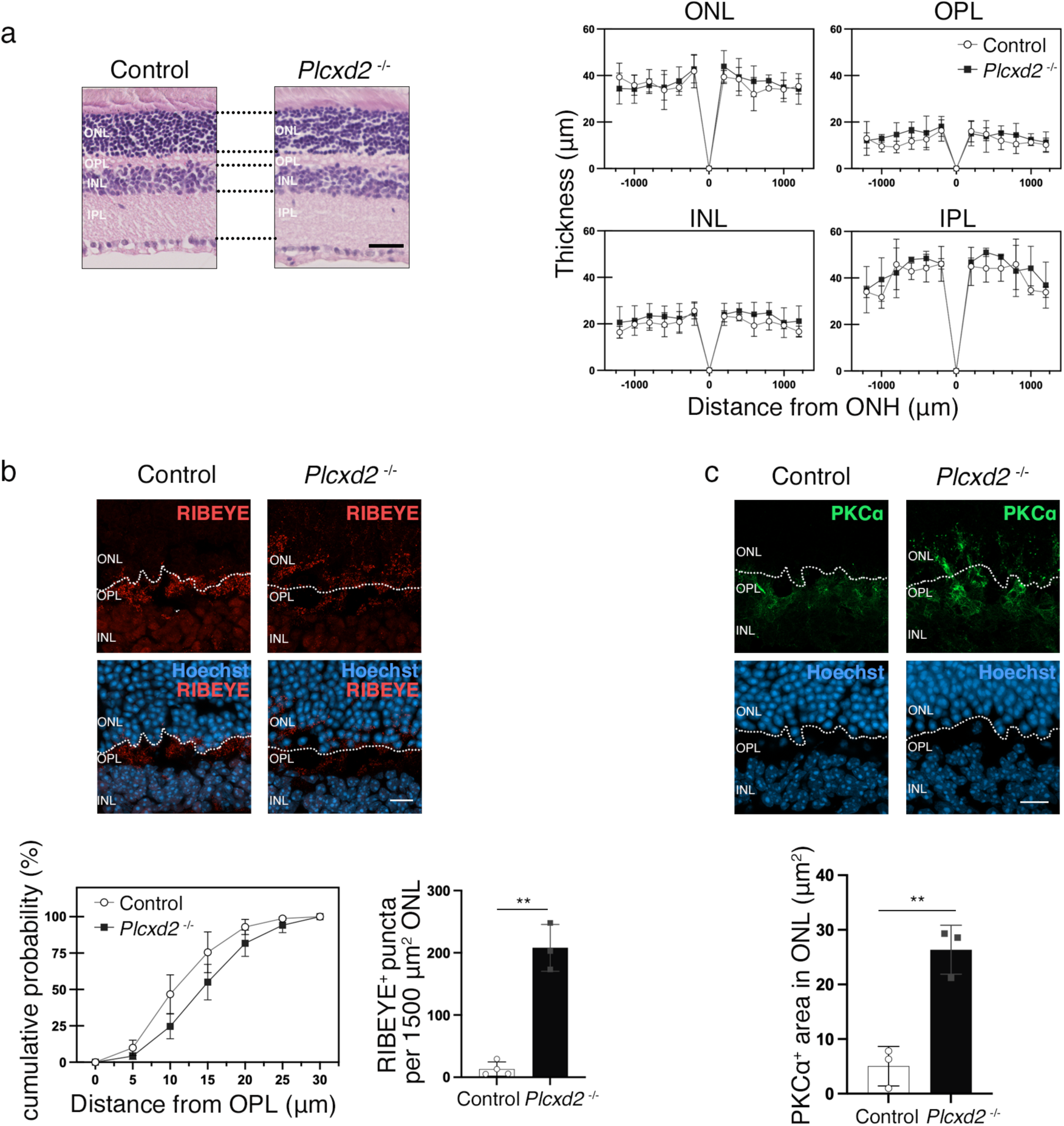
PLCXD2 deficiency causes early abnormalities at the outer retinal synaptic interface. (a) Representative hematoxylin and eosin-stained retinal sections from 3-month-old control and *Plcxd2* knockout mice and quantification of the thicknesses of the ONL, OPL, INL, and IPL relative to the optic nerve head (ONH). Scale bar, 30 µm. Control and knockout, n = 3 mice each. (b) RIBEYE immunostaining (red) with Hoechst nuclear counterstaining (blue). The cumulative distribution of RIBEYE-positive puncta was plotted as a function of their distance from the OPL toward the ONL. RIBEYE-positive puncta within a fixed 1,500-µm² region of the ONL were also quantified. Scale bar, 10 µm. Control, n = 4 mice; knockout, n = 3 mice. (c) PKCα immunostaining of rod bipolar cells (green) with Hoechst nuclear counterstaining (blue). PKCα-positive processes extending beyond the OPL into the ONL were quantified as the PKCα-positive area within the ONL. Control and knockout, n = 3 mice each. Scale bar, 10 µm. Data are presented as mean ± SD. Statistical significance was assessed using two-tailed Welch’s t-tests for the quantitative comparisons; no statistical test was applied to the cumulative distance plot. *P < 0.05, and **P < 0.01. ONL, outer nuclear layer; OPL, outer plexiform layer; INL, inner nuclear layer; IPL, inner plexiform layer.

Despite preservation of retinal layer thickness, abnormalities were already evident at the photoreceptor–bipolar synaptic interface. We first examined photoreceptor ribbon synapses using RIBEYE immunostaining. In control retinas, RIBEYE-positive structures were concentrated within the OPL, whereas in *Plcxd2* knockout retinas their distribution extended farther into the ONL (Fig. 4b). Quantification confirmed a significant increase in ectopic RIBEYE-positive puncta within the ONL in *Plcxd2* knockout retinas (Fig. 4b), indicating abnormal organization of photoreceptor presynaptic ribbons^20–23^. Consistent with remodeling of the photoreceptor–bipolar synaptic interface, PKCα-positive processes of rod bipolar cells extended beyond the OPL into the ONL in *Plcxd2* knockout retinas (Fig. 4c).

By 6 months of age, a clear structural phenotype had emerged. The ONL was significantly thinner across multiple retinal positions in *Plcxd2* knockout mice, whereas the thickness of the OPL, INL, and IPL remained largely preserved (Fig. 5a). Consistent with ongoing photoreceptor loss, the percentage of TUNEL-positive nuclei was significantly increased within the ONL of knockout retinas (Fig. 5b). Thus, the early abnormalities at the photoreceptor–bipolar synaptic interface were followed by selective degeneration of the photoreceptor-containing ONL.

**Figure 5.**
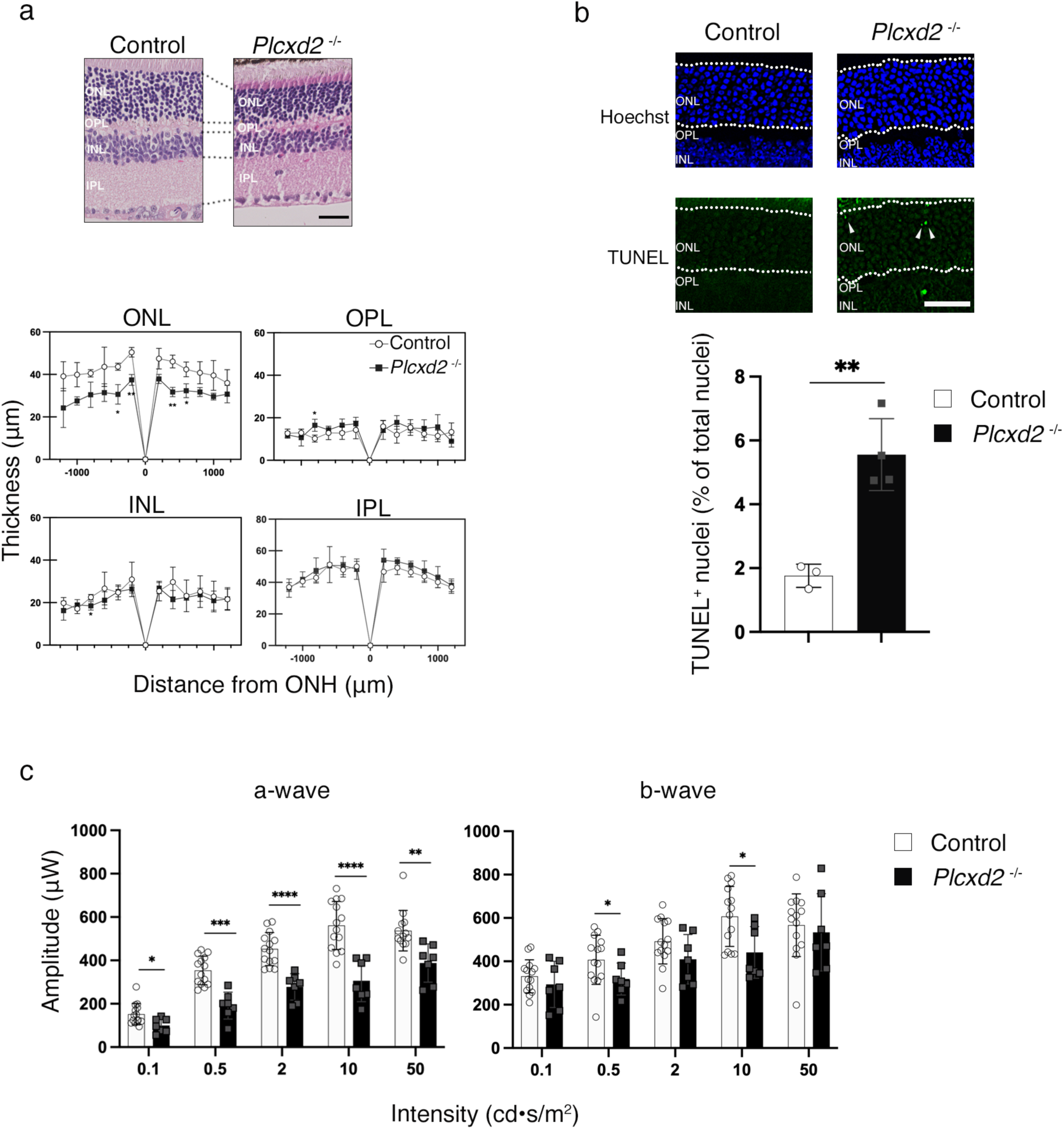
PLCXD2 deficiency leads to photoreceptor loss and impaired photoreceptor responses. (a) Representative hematoxylin and eosin-stained retinal sections from 6-month-old control and *Plcxd2* knockout mice and quantification of the thicknesses of the ONL, OPL, INL, and IPL relative to the ONH. Scale bar, 30 µm. Control and knockout, n = 3 mice each. (b) TUNEL staining of 6-month-old retinas with Hoechst nuclear counterstaining. TUNEL-positive nuclei within the ONL were quantified as the percentage of total nuclei within a fixed 1,000 µm² region. Control, n = 3 mice; knockout, n = 4 mice. (c) Scotopic electroretinography in 6-month-old control and *Plcxd2* knockout mice. A-wave and b-wave amplitudes were measured in response to flashes of 0.1, 0.5, 2, 10, and 50 cd·s/m² under dark-adapted conditions. Both male and female mice were included, and one left eye per mouse was analyzed (control, n = 13-14 mice; knockout, n = 7 mice). Data are presented as mean ± SD. Statistical significance was assessed using two-tailed Welch’s t-test. *P < 0.05, **P < 0.01, ***P < 0.001, and ****P < 0.0001. ONH, optic nerve head; ONL, outer nuclear layer; OPL, outer plexiform layer; INL, inner nuclear layer; IPL, inner plexiform layer.

We next assessed retinal function by scotopic electroretinography. A-wave amplitudes, which primarily reflect photoreceptor responses^24,25^, were markedly reduced in *Plcxd2* knockout mice across all light intensities examined (Fig. 5c). B-wave amplitudes showed more limited changes, with significant reductions detected only at 0.5 and 10 cd·s/m². Thus, the ERG phenotype was dominated by impaired photoreceptor responses, with more restricted changes in b-wave responses. This pattern was consistent with the selective ONL thinning and increased photoreceptor cell death observed histologically.

Consistent with the prominent photoreceptor phenotype, analysis of publicly available single-cell RNA-sequencing data from the Mouse Retina Cell Atlas (MRCA)^26^ showed that *Plcxd2* transcripts were detectable in both rod and cone photoreceptors, as well as in several other retinal neuronal classes (Supplementary Fig. 3).

Together, these findings reveal an age-dependent sequence of retinal abnormalities following loss of PLCXD2. Altered organization of photoreceptor ribbon synapses and associated remodeling of rod bipolar cell processes were detectable before overt retinal thinning, whereas later stages were characterized by photoreceptor cell death, loss of ONL thickness, and impaired photoreceptor responses.

## Discussion

Our study defines the substrate preference of PLCXD2, links its catalytic activity to cellular PI-linked lipid metabolism, and identifies an endogenous role for PLCXD2 in retinal lipid homeostasis and outer retinal integrity. Direct comparison under matched biochemical conditions showed that PLCXD2 hydrolyzes unphosphorylated PI substantially more efficiently than PI(4,5)P₂ and most other phosphoinositides tested. Cellular lipidomics further revealed a pronounced reduction in PI 38:4 following expression of catalytically active PLCXD2, accompanied by marked accumulation of DAG and PA. *In vivo*, loss of endogenous PLCXD2 reduced retinal DAG and PA without broadly altering major membrane phospholipid classes and was associated with progressive abnormalities of the outer retina. Together, these findings identify PLCXD2 as a PI-preferring mammalian PLC and clarify a previously unresolved distinction between PI- and PI(4,5)P₂-directed models of PLCXD2 activity.

### PI preference reframes the biochemical function of PLCXD2

Previous studies linked PLCXD2 activity to PI(4,5)P₂ depletion, but did not establish whether PI(4,5)P₂ is its preferred direct substrate^16^. Our biochemical analyses clarify this distinction. Under matched assay conditions, recombinant PLCXD2 generated substantially more DAG from PI than from PI(4,5)P₂. These endpoint assays were designed to compare substrate utilization under matched conditions rather than to define kinetic parameters. PLCXD2 also hydrolyzed PI5P, whereas no clear catalytic activity-dependent hydrolysis was detected for PI4P or most other phosphoinositides tested. Thus, PLCXD2 is not simply a conventional PI(4,5)P₂-directed PLC; rather, its substrate profile shows a pronounced preference for unphosphorylated PI. The detectable activity toward PI5P nevertheless indicates that this preference is relative rather than absolute. Because PI5P is substantially less abundant than PI in mammalian cells^10,11^, its quantitative contribution to PLCXD2-dependent lipid metabolism is likely to be limited.

This preference provides a different framework for interpreting PLCXD2-dependent changes in phosphoinositide homeostasis. Cellular PI(4,5)P₂ depletion does not necessarily imply that PI(4,5)P₂ is the predominant direct substrate. Preferential hydrolysis of the PI precursor pool could simultaneously generate DAG and alter the availability of PI for phosphoinositide synthesis, providing a potential explanation for the more modest, 38:4-centered changes observed in PIP and PIP₂. This model does not exclude physiologically relevant direct hydrolysis of PI(4,5)P₂, which was measurable *in vitro*, and substrate utilization in cells may additionally depend on membrane composition, lipid accessibility, interacting proteins, and subcellular localization.

### PI hydrolysis provides a route to both phosphoinositide and DAG–PA metabolism

PI preference has important consequences because PI occupies a different metabolic position from PI(4,5)P₂. Whereas PI(4,5)P₂ is a relatively small regulatory lipid pool, PI is the common precursor for phosphoinositide synthesis and represents a substantially larger reservoir of lipid substrate^10–12^. PLCXD2-mediated PI hydrolysis could therefore simultaneously generate DAG and alter the availability of PI for subsequent phosphorylation. The cellular lipidomic data are consistent with this substrate preference. In HEK293 cells, catalytically active PLCXD2 caused a pronounced reduction in the abundant PI 38:4 species, accompanied by a reduction in PIP 38:4 and a downward trend in PIP₂ 38:4, whereas total PIP₂ showed only a modest, statistically nonsignificant decrease. This pattern is more consistent with perturbation of the PI precursor pool than with broad depletion of PI(4,5)P₂, although steady-state cellular lipid measurements alone cannot identify the direct enzymatic substrate.

A second consequence was the pronounced increase in DAG and PA. The increase in DAG 38:4 was consistent with the decrease in PI 38:4 and with direct conversion of PI to DAG by PLCXD2. Nevertheless, changes in DAG and PA extended across multiple acyl-chain compositions, indicating that the steady-state cellular response extends beyond simple one-to-one conversion of individual PI species. DAG phosphorylation, lipid reacylation, and exchange of intermediates among interconnected glycerophospholipid pathways may contribute to this broader pattern^18,19,27^. The present experiments do not resolve these downstream reactions and therefore do not constitute a metabolic flux analysis. Nevertheless, the convergence of cellular PI loss, direct biochemical PI preference, and catalytic activity-dependent accumulation of DAG and PA provides complementary evidence that PLCXD2 can feed PI-linked lipid into the DAG–PA axis.

Importantly, the changes induced by PLCXD2 did not follow a uniform pattern across major membrane phospholipid classes. This argues against nonspecific disruption of membrane lipid composition and supports a preferential impact on PI-linked pathways. Thus, access to the abundant PI pool may allow PLCXD2 to influence cellular lipid metabolism in a manner qualitatively different from PLC activity restricted predominantly to PI(4,5)P₂: PI hydrolysis can affect both the precursor side of the phosphoinositide network and the generation of DAG available to downstream metabolic and signaling pathways.

### Endogenous PLCXD2 contributes to retinal DAG–PA homeostasis

The retinal lipidomic phenotype complements the gain-of-function and biochemical experiments by revealing the consequences of endogenous PLCXD2 loss. Retinal lipidomics was used to assess the steady-state lipid consequences of PLCXD2 deficiency in mature retina and was not designed to establish the temporal relationship between lipid changes and the earlier structural abnormalities. Loss of PLCXD2 significantly reduced retinal DAG and PA, whereas major membrane phospholipid classes did not show comparable changes in total abundance. These alterations are therefore unlikely to represent a nonspecific collapse of retinal membrane lipid composition.

Neither total PI nor PI 38:4 accumulated detectably in *Plcxd2*-deficient retina. The absence of substrate accumulation does not contradict PI utilization by PLCXD2. Steady-state abundance reflects the balance of multiple synthetic and consumptive pathways, and PI in particular is subject to extensive metabolic regulation^12^. Loss of a single route of PI consumption may consequently be accommodated without producing measurable accumulation of the bulk PI pool. Our data therefore demonstrate preservation of retinal PI abundance despite PLCXD2 deficiency but do not identify the mechanisms that maintain this steady state.

The species-level patterns also differed between the gain- and loss-of-function systems. Whereas 38:4 lipids showed the most prominent changes following PLCXD2 expression in HEK293 cells, the retinal phenotype involved changes across several DAG and PA molecular species, while the predominant DAG 38:4 species remained comparatively preserved. These differences indicate that the molecular-species pattern associated with PLCXD2 activity is strongly context dependent and should not be interpreted as evidence for a fixed preference toward 38:4 lipids. The lipid species that change most prominently are likely to reflect substrate availability, subcellular compartmentalization, and downstream metabolic processing in a given cell or tissue. Our biochemical experiments establish a preference of PLCXD2 for the PI headgroup over PI(4,5)P₂ under matched conditions, but do not define its acyl-chain selectivity.

### PLCXD2 has a physiological role in outer retinal maintenance

Loss of PLCXD2 also produced a progressive retinal phenotype. At 3 months of age, overall retinal lamination and layer thickness remained preserved, yet abnormalities were already present at the photoreceptor–bipolar synaptic interface. RIBEYE-positive photoreceptor ribbon structures showed an altered distribution toward the ONL, accompanied by ectopic extension of PKCα-positive rod bipolar cell processes into the ONL. RIBEYE is a principal structural component of retinal ribbon synapses, and perturbation of ribbon organization can be accompanied by remodeling of postsynaptic elements^20,22,28^. These changes therefore preceded overt photoreceptor loss and identify the outer retinal synaptic region as an early site affected by PLCXD2 deficiency.

By 6 months, *Plcxd2*-deficient mice exhibited selective thinning of the photoreceptor-containing ONL together with an increased proportion of TUNEL-positive nuclei and reduced scotopic ERG a-wave amplitudes. B-wave amplitudes showed more limited changes, with significant reductions detected only at selected stimulus intensities, whereas inner retinal layer thickness remained comparatively preserved. These findings indicate that the functional phenotype at this stage was dominated by impaired photoreceptor responses, although more restricted alterations in b-wave responses were also present. This functional pattern was consistent with the selective structural degeneration of the outer retina. The age-dependent pattern, with early outer retinal synaptic abnormalities followed by photoreceptor loss and impaired photoreceptor responses, supports a physiological requirement for PLCXD2 in long-term outer retinal integrity. This progressive phenotype is consistent with the broader principle that diverse molecular defects can converge on photoreceptor degeneration and subsequent retinal circuit remodeling^29,30^.

These retinal findings are also noteworthy in light of the recently described role of PLCXD2 in postsynaptic organization in cortical neurons. GPR158 restrains PLCXD2 activity to regulate spine apparatus incorporation and dendritic spine maturation^16^, whereas the present study identifies early abnormalities at another specialized neuronal synaptic interface following PLCXD2 loss. Together, these observations suggest that appropriate regulation of PLCXD2 may be particularly important in highly specialized neuronal membrane environments. Whether these phenotypes share a common dependence on PLCXD2-mediated PI metabolism remains to be established.

Beyond the nervous system, PLCXD2 has also been implicated in human disease. Elevated PLCXD2 expression was recently reported in head and neck squamous cell carcinoma, where higher expression was associated with advanced tumor stage and poorer patient outcome^31^. Whether PLCXD2 catalytic activity or altered PI-linked lipid metabolism contributes mechanistically to these cancer-associated phenotypes remains unknown. The substrate preference defined here provides a biochemical basis for addressing this question.

An important limitation is that our data do not establish a causal link between the retinal DAG–PA changes and the structural and functional retinal phenotypes. These findings therefore identify parallel consequences of PLCXD2 deficiency, while the mechanistic relationship between them remains unresolved. Conversely, the selective nature of the retinal lipid changes suggests that they are unlikely to reflect solely a nonspecific consequence of retinal degeneration.

Together, our findings establish PLCXD2 as a PI-preferring mammalian PLC and reveal an endogenous role for PLCXD2 in retinal DAG–PA homeostasis and long-term outer retinal integrity. More broadly, they expand the functional repertoire of mammalian PLCs beyond canonical PI(4,5)P₂-directed signaling and provide a framework for understanding the physiological roles of atypical PLCXD proteins.

## Methods

### Lipids and reagents

Synthetic PI, phosphoinositides, phospholipids, and lipid standards were obtained from Avanti Polar Lipids (Merck) unless otherwise stated. DAG 17:1/17:1 (diheptadecenoin; Nu-Chek Prep, D-405) was used as an internal standard for DAG in retinal lipidomics. HPLC- or MS-grade solvents were obtained from Fujifilm Wako Pure Chemical, trimethylsilyldiazomethane from Tokyo Chemical Industry, and DEAE Sepharose Fast Flow from Cytiva.

### Cell culture, transfection and immunoblotting

HEK293 cells were maintained in Dulbecco’s modified Eagle’s medium containing 1.0 g/L glucose, L-glutamine, and sodium pyruvate (Nacalai Tesque), supplemented with 10% fetal bovine serum (Nichirei Biosciences) and 1% penicillin–streptomycin (Nacalai Tesque), at 37 °C in a humidified atmosphere containing 5% CO₂. Cells were transiently transfected with empty pCAGGS vector or pCAGGS plasmids encoding FLAG-tagged human wild-type PLCXD2 (PLCXD2 WT) or the previously characterized catalytically inactive H57L/H132L mutant (PLCXD2 MT) using polyethylenimine (Polysciences). Cells were harvested 24 h after transfection for lipidomic analysis. PLCXD2 expression was confirmed by SDS–PAGE and immunoblotting using anti-FLAG antibody (F1804, Sigma-Aldrich; 1:1,000). GAPDH (sc-365062, Santa Cruz Biotechnology; 1:1,000) was used as a loading control.

### Cellular lipidomics

Cell pellets were suspended in formic acid, and class-appropriate internal standards were added before lipid extraction. PI and phosphoinositides were extracted under acidic conditions, enriched by DEAE chromatography, and methylated with trimethylsilyldiazomethane essentially as described previously^32,33^, with modifications appropriate for the present analysis. The following internal standards were used: PI 15:0/18:1-d7 for PI; PI4P 17:0/20:4 for PIP; PI(4,5)P2 17:0/20:4 for PIP2. Reverse-phase LC–MS/MS analysis of PI and phosphoinositides was performed in positive-ion multiple-reaction-monitoring (MRM) mode using a Nexera XR system (Shimadzu) coupled to an LCMS-8050RX triple-quadrupole mass spectrometer (Shimadzu). Lipids were separated on a COSMOCORE 2.6C18 column (2.1 × 150 mm, 2.6 µm; Nacalai Tesque) maintained at 60 °C. Mobile phase A consisted of acetonitrile/water/ethylamine (240:60:0.195, v/v/v), and mobile phase B consisted of isopropanol/acetonitrile/ethylamine (240:60:0.195, v/v/v). The gradient was as follows: 0–1 min, 30% B; 1–3 min, a linear increase from 30% to 90% B; 3–7.5 min, 90% B; and 7.5–12 min, 30% B. The flow rate was 220 µL/min. Peak areas were normalized to those of the corresponding internal standards. For analyses of DAG, PA, and major membrane phospholipids, total lipids were extracted from cells using the method of Bligh and Dyer^34^. DAG was analyzed in positive-ion MRM mode using a Nexera X2 system (Shimadzu) coupled to a QTRAP 7500 triple-quadrupole mass spectrometer (SCIEX). Lipids were separated on an ACQUITY UPLC HSS T3 column (2.1 × 100 mm, 1.8 µm; Waters) using chromatographic conditions adapted from a previously described method^35^. DAG levels are presented as LC–MS/MS peak areas normalized to 1 × 10^5^ cells. PA and other major membrane phospholipids were analyzed in negative-ion MRM mode using a Nexera X2 system (Shimadzu) coupled to a QTRAP 4500 triple-quadrupole mass spectrometer (SCIEX). Lipids were separated on a SeQuant ZIC-HILIC PEEK-coated column (2.1 × 250 mm, 1.8 µm; Merck Millipore) using chromatographic conditions adapted from a previously described method^36^. Peak areas were normalized to those of the corresponding internal standards.

### Recombinant protein preparation and *in vitro* PLC assay

FLAG-tagged human PLCXD2 WT and PLCXD2 MT, together with human PLCδ1 and *Listeria monocytogenes* PI-PLC, were produced using a wheat-germ cell-free proteoliposome expression system (CellFree Sciences). Protein preparations were assessed by SDS–PAGE followed by Coomassie Brilliant Blue staining. For phospholipase assays, PC 32:0 and the indicated PI or phosphoinositide substrate [PI, PI3P, PI4P, PI5P, PI(3,4)P₂, PI(3,5)P₂, PI(4,5)P₂, or PI(3,4,5)P₃] with an 18:0/20:4 acyl-chain composition were mixed at final concentrations of 9 and 5 μM, respectively. The lipids were dried under a stream of nitrogen and resuspended by sonication and vortexing in 15 μL of assay buffer containing 50 mM HEPES–NaOH (pH 6.8), 100 mM NaCl, 3 mM CaCl₂, and 2.5 mM EGTA. Protein preparations (3 μg in 5 μL of assay buffer) were added to the lipid suspension to give a final reaction volume of 20 μL. Reactions were incubated at 30 °C for 15 min. Reactions lacking recombinant protein preparation were used as no-enzyme controls. Reactions were terminated by addition of chloroform/methanol/HCl/water (120:120:10:10, v/v/v/v), and DAG production was quantified by LC–MS/MS as a measure of PLC activity. LC–MS/MS analysis of DAG was performed in positive-ion MRM mode using a SCIEX 5500 triple-quadrupole mass spectrometer. Lipids were separated on a COSMOCORE 2.6C18 column (2.1 × 150 mm, 2.6 µm; Nacalai Tesque) maintained at 60 °C. Mobile phase A consisted of isopropanol/acetonitrile/1 M ammonium acetate (160:40:1, v/v/v), and mobile phase B consisted of acetonitrile/ultrapure water/1 M ammonium acetate (160:40:1, v/v/v). The gradient was as follows: 0–1 min, 30% A; 1–3 min, a linear increase from 30% to 90% A; 3–7.5 min, 90% A; and 7.5–12 min, 30% A. The flow rate was 220 µL/min. For quantification of DAG 18:0/20:4 (DAG 38:4), the ammonium adduct [M+NH₄]⁺ at m/z 662.572 was selected in Q1 and subjected to collision-induced dissociation. The product ion corresponding to 18:0 monoacylglycerol (MAG 18:0) at m/z 341.305 was monitored, giving an MRM transition of m/z 662.572 → 341.305. The declustering potential, entrance potential, collision energy, and collision-cell exit potential were 100, 10, 30, and 12 V, respectively. DAG 15:0/18:1-d7 was used as an internal standard. Its ammonium adduct at m/z 605.584 and the product ion corresponding to 15:0 MAG at m/z 299.258 were monitored using the MRM transition m/z 605.584 → 299.258, with a collision energy of 28 V. DAG production was quantified by normalizing the peak area of the 18:0 MAG product ion to that of the 15:0 MAG product ion derived from the internal standard.

### Generation and maintenance of *Plcxd2* knockout mice

*Plcxd2* knockout mice were generated by Cyagen using CRISPR/Cas9-mediated genome editing. Two guide RNAs targeting regions flanking exons 2 and 3 were used to generate a deletion encompassing both exons. Mice were maintained on a C57BL/6N background under specific-pathogen-free conditions on a 12-h light/12-h dark cycle with ad libitum access to standard chow and water. Male mice were used for all experiments except electroretinography, for which both male and female mice were used. Wild-type and heterozygous littermates were used as controls. All procedures were approved by the Animal Experimentation Committee of Tokyo University of Science and were performed in accordance with institutional guidelines.

### Genotyping and retinal cDNA analysis

Genomic DNA prepared from tail biopsies was analyzed by PCR using Quick Taq HS DyeMix (TOYOBO). The F1, R1, and R2 primers used to distinguish the wild-type and deletion alleles were 5′-CCCCGCTAAGACTGAAGAAACTAC-3′, 5′-CACCCAGCTGTCTCTACTTCTCTA-3′, and 5′-TCTTATTAGCGGCTCTTTAAGCCT-3′, respectively. PCR was performed for 35 cycles of 94 °C for 30 s, 60 °C for 30 s, and 68 °C for 30 s. Total retinal RNA was isolated using a FastGene RNA kit (Nippon Genetics) and reverse-transcribed using ReverTra Ace qPCR RT Master Mix with gDNA Remover (TOYOBO). *Plcxd2* cDNA spanning the deleted region was amplified using the primers 5′-ATGCTTGCATTTAGAAAGGC-3′ and 5′-TCAACACTTAGTCAGAGCTC-3′. PCR products were analyzed by Sanger sequencing.

### Histology and retinal layer measurements

Eyes from 3- and 6-month-old mice were fixed in 4% paraformaldehyde in PBS, embedded in paraffin, and sectioned at 5 µm. Sections were stained with Mayer’s hematoxylin and eosin Y and imaged using a BZ-X810 microscope (Keyence). The thicknesses of the ONL, OPL, INL, and IPL were measured at predefined distances from the optic nerve head on both sides of the retina using ImageJ (NIH).

### Immunofluorescence and image analysis

Eyes were embedded in O.C.T. Compound (Sakura Finetek Japan), frozen, and sectioned at 10 µm. Sections were fixed in 4% paraformaldehyde in PBS, permeabilized and blocked in PBS containing 3% BSA and 0.1% Triton X-100, and incubated overnight at 4 °C with anti-RIBEYE (192003, Synaptic Systems; 1:1,000) or anti-PKCα (sc-8393, Santa Cruz Biotechnology; 1:200). Alexa Fluor-conjugated secondary antibodies (Thermo Fisher Scientific) were used for detection, and nuclei were counterstained with Hoechst 33342 (Nacalai Tesque). Images were acquired using a BZ-X810 microscope or an LSM 900 confocal microscope (Carl Zeiss). At least three images per animal were analyzed using ImageJ, and values from multiple images from the same animal were averaged before statistical analysis. For RIBEYE analysis, RIBEYE-positive puncta were identified within defined regions of the outer retina. Their positions were quantified according to the distance from the OPL boundary toward the ONL and used to generate cumulative distance distributions. In addition, the number of RIBEYE-positive puncta within a fixed 1,500-μm² region of the ONL was quantified. For PKCα analysis, the PKCα-positive area within a fixed 1,500-μm² region of the ONL was quantified.

### TUNEL assay

Apoptotic cells in retinas from 6-month-old mice were detected in 5-µm paraffin sections using the In Situ Apoptosis Detection Kit (TaKaRa Bio). Following deparaffinization, rehydration, and proteinase K treatment, sections were incubated with labeling solution and rTdT enzyme for 80 min at 37 °C according to the manufacturer’s instructions. Images were acquired using a BZ-X810 microscope. TUNEL-positive and total Hoechst-positive nuclei were counted within a fixed 1,000 µm² region of the ONL using ImageJ, and the percentage of TUNEL-positive nuclei among total nuclei was calculated.

### Retinal lipidomics

Retinas were collected from control and *Plcxd2* knockout littermates at 21 weeks of age. Retinal lipids were extracted using the method of Bligh and Dyer^34^ with class-specific internal standards. The following internal standards were used: PC 15:0/15:0 and PC 15:0–18:1-d7 for PC; PE 15:0/15:0 and PE 15:0–18:1-d7 for PE; PG 14:0/14:0 for PG; PS 14:0/14:0 for PS; PI 15:0/18:1 for PI; DAG 15:0/18:1-d7 and DAG 17:1/17:1 for DAG; PA 17:0/17:0 for PA; and SM 16:0-d31 for SM. The organic phase was transferred to a clean vial and dried under a stream of nitrogen. Lipids were reconstituted in methanol and stored at −80 °C until analysis. A portion of each lipid extract was analyzed by ultrahigh-performance liquid chromatography–electrospray ionization tandem mass spectrometry (LC–ESI–MS/MS). For quantification of PA and PS, LC separation was performed on an ACQUITY Premier BEH C18 column (1.7 µm, 2.1 × 50 mm; Waters). Mobile phase A consisted of H₂O/acetonitrile (80:20, v/v) containing 0.028% NH₄OH, and mobile phase B consisted of isopropanol/acetonitrile (80:20, v/v). The LC method consisted of a linear gradient from 100% A to 100% B over 17.5 min, 100% B for 5 min, a linear gradient to 100% A over 2.5 min, and equilibration with 100% A for 5 min, for a total run time of 30 min. The flow rate was 0.2 mL/min, and the column temperature was 25 °C. For quantification of the other lipid classes, LC separation was performed on an ACQUITY UPLC BEH C18 column (1.7 µm, 2.1 × 100 mm; Waters) coupled to an ACQUITY UPLC BEH C18 VanGuard pre-column (1.7 µm, 2.1 × 5 mm; Waters). Mobile phase A consisted of acetonitrile/water (60:40, v/v) containing 10 mM ammonium formate and 0.1% (v/v) formic acid, and mobile phase B consisted of isopropanol/acetonitrile (90:10, v/v) containing 10 mM ammonium formate and 0.1% (v/v) formic acid. The LC gradient consisted of 20% B for 2 min, a linear increase to 60% B over 4 min, a linear increase to 100% B over 16 min, and equilibration with 20% B for 5 min, for a total run time of 27 min. The flow rate was 0.3 mL/min, and the column temperature was 55 °C. MRM analysis was performed using a Xevo TQ-S micro triple-quadrupole mass spectrometer (Waters) equipped with an ESI source. The ESI capillary voltage was set to 1.0 kV, and the sampling cone voltage was 30 V. The source and desolvation temperatures were 150 °C and 500 °C, respectively. The desolvation and cone gas flow rates were 1,000 and 50 L/h, respectively. Lipid signals were normalized to the corresponding class-specific internal standards. Because the small size of the retina made normalization to tissue weight unreliable, internal-standard-normalized lipid abundance was further normalized to total PC measured in each sample.

### Electroretinography

Scotopic electroretinograms (ERGs) were recorded using a Ganzfeld dome, an acquisition system (PuREC, MAYO Corporation) and an LED stimulator. Following overnight dark adaptation, mice were anesthetized under dim red light with a mixture of medetomidine hydrochloride (0.75 mg/kg; Nippon Zenyaku Kogyo), midazolam (4 mg/kg; Sandoz) and butorphanol tartrate (5 mg/kg; Meiji Seika Pharma) (MMB). Pupils were dilated with tropicamide and phenylephrine hydrochloride (Santen Pharmaceutical). Body temperature was maintained using a heating pad throughout the recordings. Corneal contact electrodes were used as active electrodes, a reference electrode was placed in the mouth, and a clip electrode attached to the tail served as the ground. ERG responses were recorded under dark-adapted conditions using white-flash stimuli of 0.1, 0.5, 2, 10 and 50 cd·s/m². Signals were processed post-hoc using a 300-Hz low-pass filter for a-wave analysis and a 30-Hz low-pass filter for b-wave analysis.

### Analysis of public single-cell RNA-sequencing data

Publicly available single-cell RNA-sequencing data from the Mouse Retina Cell Atlas (MRCA)^26^ were analyzed. The integrated dataset release analyzed here comprised 323,957 cells assigned to 11 major retinal cell classes. Cells from all available ages were included and pooled for analysis. Cells with a raw *Plcxd2* UMI count greater than zero were defined as *Plcxd2*-positive, and the percentage of positive cells was calculated for each major retinal cell class as the number of *Plcxd2*-positive cells divided by the total number of cells in that class. To provide an additional descriptive measure of transcript abundance among cells in which *Plcxd2* was detected, the mean log1p-transformed counts per 10,000 transcripts [log1p(CP10K)] was calculated among *Plcxd2*-positive cells for which normalized CP10K values were available. Because normalized expression values were not available for all cells in the integrated dataset, this measure was used descriptively and was not subjected to statistical comparison between retinal cell classes.

### Statistics and Reproducibility

Data are presented as mean ± SD unless otherwise indicated. For image-based analyses, measurements from multiple fields or sections were averaged to obtain one value per animal before statistical testing. Two-group comparisons were performed using two-tailed Welch’s t-tests. Comparisons among three or more groups were performed using one-way ANOVA followed by Tukey’s or Dunnett’s multiple-comparisons test, as specified in the figure legends. The biological sample size and definition of independent replicates for each experiment are indicated in the figure legends. P < 0.05 was considered statistically significant.

## Acknowledgements

We thank K. Sundquist, M. Yashiro, M. Morita, H. Kuroda, A. Mizutani, and R. Usami (Tokyo University of Science) for technical assistance. This work was supported by the Japan Society for the Promotion of Science (JSPS) Grant-in-Aid for Challenging Research (Exploratory) (26K23204), JSPS Grant-in-Aid for Scientific Research (C) (23K06103), JSPS Grant-in-Aid for Early-Career Scientists (21K15109), the Nakajima Foundation, Takeda Science Foundation, The Naito Foundation, ONO Medical Research Foundation, and the Saburo Kakiuchi Memorial Award for Encouragement of Research to K.K.; and JSPS Grant-in-Aid for Scientific Research (B) (26K02823 and 23K28031), AMED-CREST from the Japan Agency for Medical Research and Development (AMED) (26gm1710007s0304), and Takeda Science Foundation to Y.N.

## Author contributions

K.K.: Conceptualization; Formal analysis; Funding acquisition; Validation; Data curation; Investigation; Visualization; Methodology; Writing—original draft; Writing—review and editing. W.I.: Data curation; Investigation. A.K.: Data curation; Investigation. F.O.: Methodology; Data curation; Investigation. Y.K.: Methodology; Data curation; Investigation. N.K.: Methodology; Data curation; Investigation; Writing—review and editing. H.-C.L.-O.: Methodology; Data curation; Investigation; Writing—review and editing. H.O.: Methodology; Data curation; Investigation; Writing—review and editing. H.K.: Investigation; Writing—review and editing. J.H.: Investigation; Writing—review and editing. S.M.: Investigation; Writing—review and editing. J.S.: Investigation; Writing—review and editing. A.H.: Investigation; Writing—review and editing. N.B.: Investigation; Writing—review and editing. J.A.: Investigation; Writing—review and editing. T.Y.: Investigation; Writing—review and editing. T.S.: Investigation; Writing—review and editing. Y.N.: Conceptualization; Formal analysis; Supervision; Funding acquisition; Investigation; Visualization; Methodology; Writing—original draft; Project administration; Writing—review and editing.

## Competing interests

The authors declare no competing interests.

---

**Supplementary Fig. 1.**
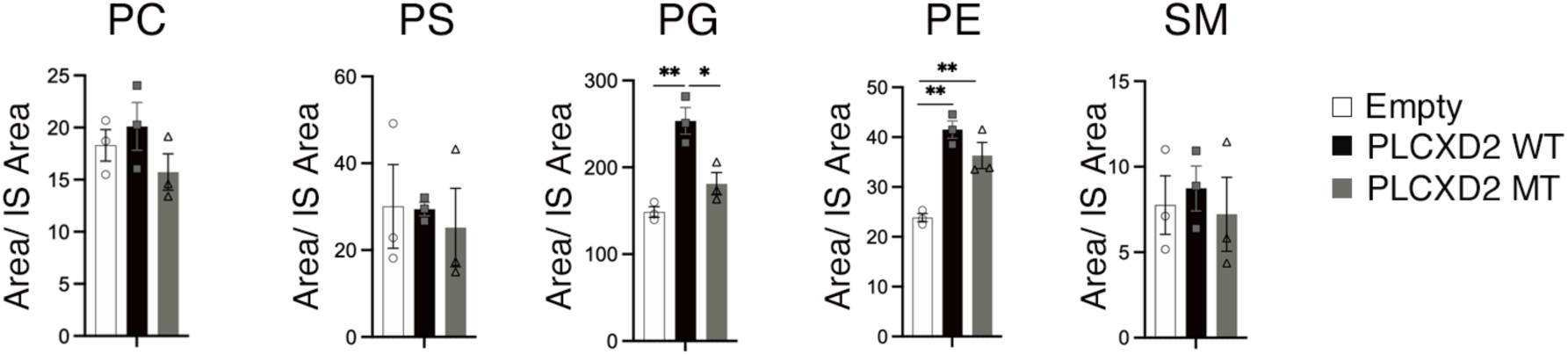
Major membrane phospholipid classes in PLCXD2-expressing HEK293 cells. LC–MS/MS analysis of PC, PS, PG, PE, and SM extracted from HEK293 cells expressing empty vector, PLCXD2 WT, or PLCXD2 MT. The summed normalized signal for all detected molecular species within each lipid class is shown. Lipid levels were calculated as the analyte peak area divided by the corresponding internal-standard (IS) peak area. Data are presented as mean ± SD from three biologically independent experiments (n = 3). Statistical significance was assessed by one-way ANOVA followed by Tukey’s multiple-comparisons test. *P < 0.05 and **P < 0.01.

**Supplementary Fig. 2.**
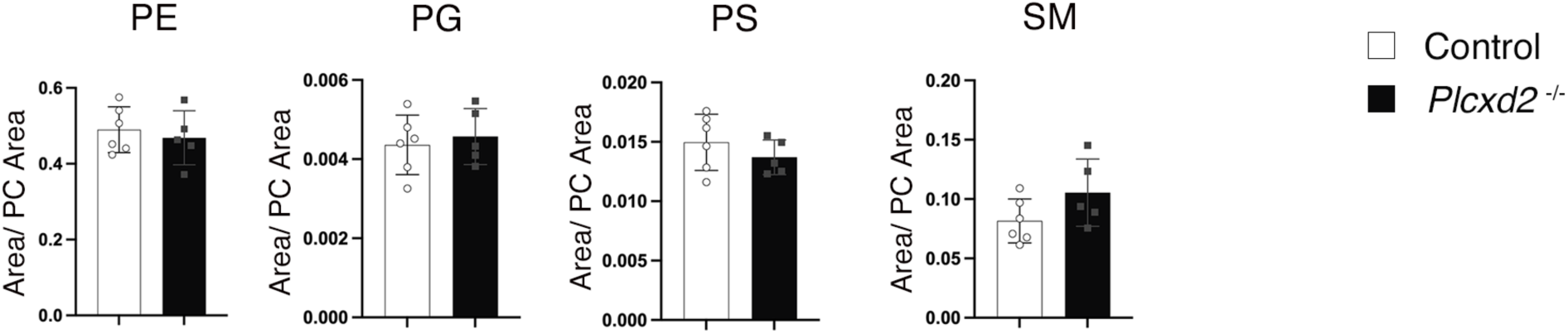
Major membrane phospholipid classes in Plcxd2-deficient retina. LC–MS/MS analysis of PE, PG, PS, and SM in retinas from 21-week-old male control and Plcxd2 knockout mice. The summed signal for all detected molecular species within each lipid class is shown. Lipid abundance was normalized to total PC within each sample. Data are presented as mean ± SD (control, n = 6 mice; Plcxd2 knockout, n = 5 mice). Statistical significance was assessed using a two-tailed Welch’s t-test.

**Supplementary Fig. 3.**
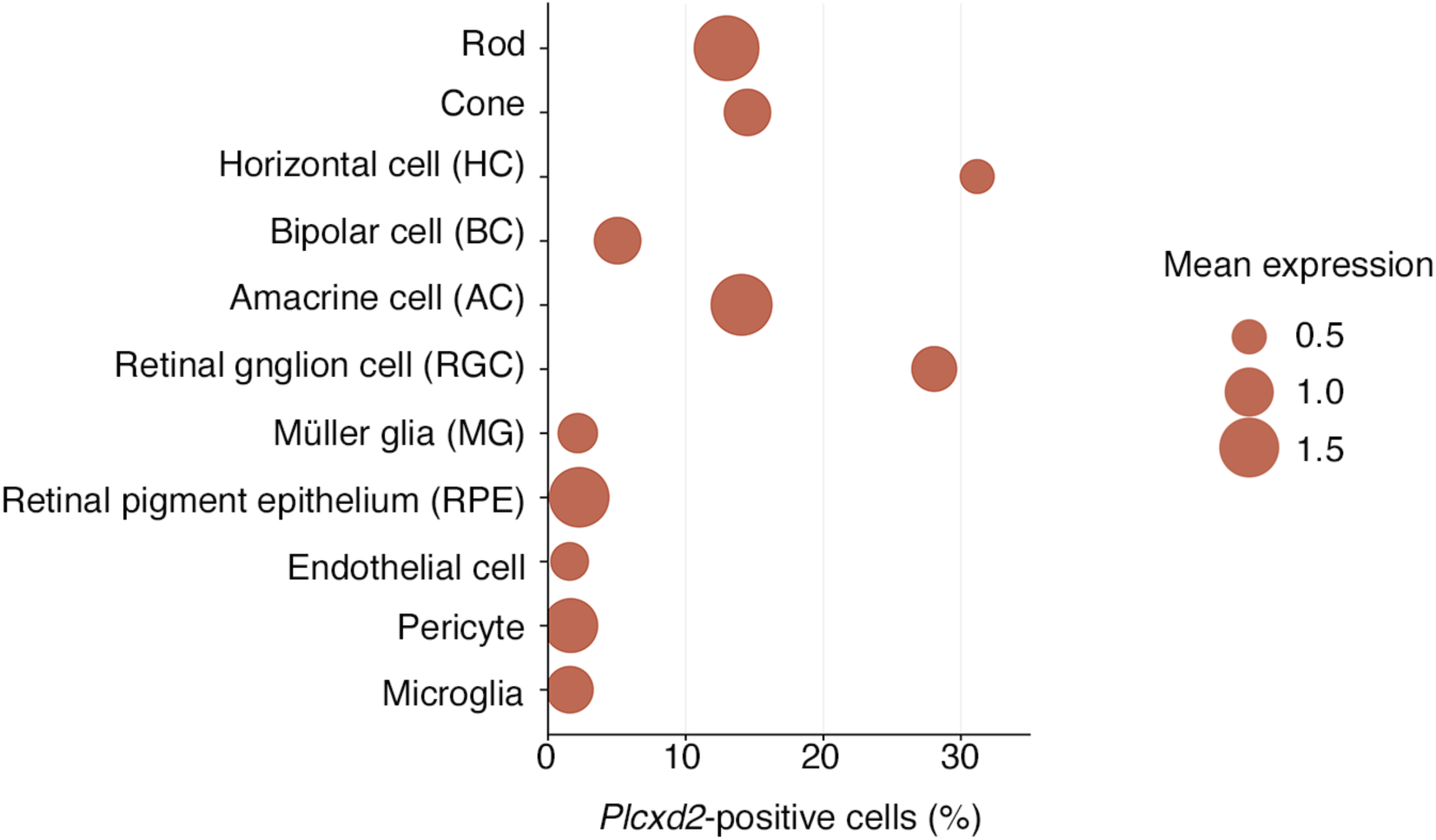
Distribution of Plcxd2 transcript detection across major retinal cell classes. Publicly available single-cell RNA-sequencing data from the Mouse Retina Cell Atlas (MRCA)^26^ were analyzed. The dataset analyzed here comprised 323,957 cells assigned to 11 major retinal cell classes, with cells from all available ages pooled for analysis. Cells with a raw *Plcxd2* UMI count greater than zero were defined as *Plcxd2*-positive. Dot position indicates the percentage of *Plcxd2*-positive cells within each major retinal cell class, whereas dot size represents the mean log1p(CP10K) value among *Plcxd2*-positive cells for which normalized expression values were available.

## Notes

### Competing Interest Statement

The authors have declared no competing interest.

